# Multiscale modeling of the spatial structure of stem cells in neuroblastoma patient-derived tumoroids reveals a critical role for a short range diffusive process

**DOI:** 10.1101/2025.07.24.666524

**Authors:** Thi Nhu Thao Nguyen, Catherine Koering, Elodie Vallin, Sandrine Gonin-Giraud, Laura Broutier, Samuel Bernard, Fabien Crauste, Olivier Gandrillon

**Author notes:** Joint last co-authors (Fabien Crauste, Olivier Gandrillon).

## Abstract

Neuroblastomas are heterogeneous pediatric tumors of the sympathetic nervous system for which treatments are still limited. Fundamental and applied approaches have been enabled thanks to the generation of patient-derived tumoroids (PDTs), *ex vivo* 3D structures used as avatars of the original tumor.

We generated neuroblastoma PDTs and quantified the spatial distribution of CD133^+^ cancer stem cells using immunohistochemistry. We observed that those cells tend to aggregate in the PDT.

In order to better understand the set of rules needed for generating such structures, we implemented a multiscale agent-based neuroblastoma tumoroid model. Model rules specify single cell’s fate based on its intracellular content, which dynamically evolves according to a stochastic gene regulatory network. The state of this network can be modulated by cell-to-cell signalling through neighbor cells fate decisions and, possibly, spatial location.

We first observed that in the absence of any spatial rules for inter-cellular interactions, no spatial structure emerged. The addition of simple rules (signalling by cell-to-cell contact or differential cell adhesion) only marginally improved the quantitative agreement to the experimental dataset. In sharp contrast, the addition of short-range pro-stem cell diffusive signalling among stem cells produced very realistic 3D PDT-like structures.

This works highlights the power of our multiscale approach to discard too simplistic rules and to propose a minimal set of hypotheses required to reproduce qualitatively and quantitatively experimentally observed spatial structures. In the case of neuroblastomas-derived PDTs, short-range spatial diffusion of stem-to-stem cell signalling proved to play a key role in successfully reconstructing the spatial structure.

## 1 Introduction

Neuroblastomas (NB) are pediatric tumors of the sympathetic nervous system derived from primitive neural crest cells. Although rare, they are the most frequent solid tumors in children under 5 years old and account for around 15% of cancer-related deaths [27]. NB is a tumor with heterogeneous biological, morphological, genetic and clinical characteristics. Treatments are still limited to a few actionable mutations [34] and very few drugs have been validated clinically [1].

The recent possibility to generate *ex vivo* relevant 3D structures that recapitulate at least part of the histological architecture typical of NB has paved the way for both fundamental and applied approaches on Patients-Derived Tumoroïds (PDTs) from children suffering from NB [17, 28]. To characterize neuroblastoma spatial structure, we developed PDTs cultures from a NB tumor and used immunohistochemistry (IHC) to assess its neuroblastoma nature. We observed a characteristic spatial positioning of CD133^+^ cancer stem cells (CSCs) that tend to be clustered around the center of the PDT. To better understand the mechanisms at work behind structure formation, we implemented a computational model of PDT growth and assessed the ability of this model to investigate and reproduce growth and differentiation spatial patterns.

Ideally the computational modelling framework should be able to accommodate the fact that cell decision making is a multiscale dynamical process, which associates an intracellular genome-based gene regulatory network (GRN), which is “stochastic at every level of organization” [7], a cellular behavior, and a tissue-level organization. Each level can both influence and be influenced by other levels. For example, it has been shown that tissue organization exerts stabilizing constraints against noise on the cellular level [21]. The cellular behavior emerges from the GRN dynamics but the molecular level can also be influenced by cellular level constraints like the cell metabolic state [36].

There is a large body of literature on stem cell modelling (whether cancerous or not) and cancer cells (whether stem or not) but only a handful of models have been used to explore the CSCs dynamics within tumors [3, 6, 12, 13, 14, 15, 16, 24, 42].

While some studies rely upon ordinary differential equations to model CSC dynamics (and therefore neglect spatial aspects; [42]), most of these studies rely upon the use of agent-based models (ABMs). All of those ABMs models are built on a 2D lattice grid, except for [6] which uses a 2D continuous space (implemented in NetLogo). Although none of the models simulate GRNs, some studies [6, 16] can be described as multiscale models: Biava et al. [6] models how molecular factors (peptides, microRNAs) influence tumor cells behaviour, connecting the molecular and cellular levels through probabilistic rules, while Fotinos et al. [16] links cell-level behavior (division, differentiation) to tissue-level patterns of CSC organization.

Additionally, while several models [3, 13, 14, 15, 24] are purely conceptual and theory-driven, demonstrating how tumor-like behavior can arise from basic cell-level rules, others (notably [6] and [16]) incorporate quantitative validation using experimental data, such as fluorescence imaging, cell marker intensity, and graph-based spatial statistics.

Reviewed studies offer valuable conceptual and computational insights into CSC dynamics and tumor growth, but they share several notable limitations. Most of these models are confined to 2D lattice-based environments, which restricts spatial realism and may oversimplify the complexity of *in vivo* tumor structure. Gene regulatory networks and intracellular signalling pathways are absent, limiting the models’ ability to capture the molecular mechanisms underlying cell fate decisions. This might prove especially limiting when the model’s output might be confronted to spatial transcriptomics data [44].

To overcome these limitations, we developed `Simuscale` [5, 32], a multiscale agent-based modeling framework that incorporates a stochastic gene regulatory network in each cell, the state of which can be influenced by cell-to-cell signalling. The resulting model is truly multiscale in the sense that it harbours at least two nested organization scales and interactions between and among scales. This results in a circular causality scheme [33] that can not be easily apprehended by more traditional modeling schemes.

`Simuscale` enables simulations from molecular to population scales, while considering both spatial movement and signaling dynamics, whether through direct cell-to-cell (local) contact or through long-range diffusion.

The model introduced in this paper consists of 15 initiating stem cells, endowed with a GRN consisting of a simple toggle-switch between two proteins controlling cell fate, and regulating a protein controlling cell proliferation. This GRN was modelled as a piecewise deterministic Markov process (PDMP) operating under the bursty regime [22], which we previously demonstrated to be able to reproduce realistic single-cell transcriptomics data [45]. In the absence of any spatial rule, the model proved insufficient to reproduce qualitatively or quantitatively the observed IHC images. The addition of simple spatial rules, like stem-to-stem cell contact and cell movement, only marginally improved thsimulation outcomes. The key to successful reconstruction of the observed spatial structure of PDTs proved to be a stem-to-stem short range diffusive signalling process.

## 2 Material, Methods and Models

### 2.1 Generating PDX

Frozen cells from Orthotopic Patient-derived xenograft (OPDX; [41]) SJNBL013762 X1 were provided by the St. Jude Children’s Research Hospital. This OPDX was derived from a tumour removed from a 15 month-old boy, with adrenal primary tumor location and bone marrow metastasis, which harboured known molecular alterations consisting in *nmyc* gene amplification and *alk* mutation.

Patient-derived xenograft (PDX) were generated according to [41]: 10^6^ thawed OPDX cells were injected subcutaneously after embedding in 100 µL matrigel (Corning™ Matrice Matrigel™ ref 356234) in 6 weeks old NSG-NOD.Cg-Prkdcscid Il2rgtm1Wjl/SzJ (Charles River ref. 614) female mice. The mice were housed in sterilized filter-topped cages and maintained in the P-PAC pathogen-free animal facility (D 69 388 0202; Cancer Research Center of Lyon; CRCL). All animal studies were performed in strict compliance with relevant guidelines validated by the local Animal Ethic Evaluation Committee (C2EA-15) and authorized by the French Ministry of Education and Research (Authorization APAFIS#28836). The study had all necessary regulatory approvals and informed consents are available for all patients.

### 2.2 Derivation and culture of neuroblastoma PDTs

Fresh tissues from tumor developed in NSG-NOD mice were mechanically minced into small pieces and enzymatic dissociation was done according to the St. Jude Children’s Research Hospital guidelines using accutase for 1-2h at 37°C. 4.10^5^ to 1.10^6^ cells were seeded after embedding in 20 µL matrigel and then placed in 100 µL medium in 96-well ULA plates (BIOFLOAT™ 96-well U bottom - FaCellitate) in a culture medium based on previously described neuroblastoma tumor-initiating cell lines and PDTs derivation protocols [4, 28] which consists of advanced DMEM/F-12 medium (Gibco), 1X B-27 supplement without vitamin A (Gibco), 1X N2 supplement (Gibco), 1x Penicillin-Streptomycin (Gibco), 40 ng*/*mL hFGF-b (Peprotech), 20 ng*/*mL hEGF (Peprotech), 10 ng*/*mL hPDGF-AA (Peprotech), 200 ng*/*mL hIGF1 (Peprotech) and 10 ng*/*mL hPDGF-BB (Peprotech).

Cultures were then split without embedding and grown in the same medium and the same 96-well ULA plates. Cultures were supplemented after 7 days by the addition of 50 µL of medium, and then the medium was renewed by half twice a week. PDTs were split every 15-19 days when reaching a diameter of about 1 mm using TrypLE Express Enzyme (ThermoFisher Scientific).

### 2.3 Histological analyses

Neuroblastoma organoids were fixed and processed as described in [19] when they reached a diameter of 0.8 mm. They were embedded in paraffin (ASP6025, Leica) and processed for histological features by the Research pathology platform East (Anapath Recherche, Cancer Research Center of Lyon (CRCL), Lyon, France). Three µm-thick tissue sections of formalin-fixed, paraffin-embedded tissue were prepared according to conventional procedures. Sections were then stained with hematoxylin and eosin and examined with a light microscope.

Immunohistochemistry was performed on an automated immunostainer (Ventana Discovery XT, Roche, Meylan, France) using Omnimap DAB Kit according to the manufacturer’s instructions. Slides were deparaffinized at 72°C using Ventana EZ Prep reagent (Ref: 950-102, Roche) and hydrated, followed by an antigen retrieval method using Ventana Tris-EDTA buffer pH 7.8 (Ref: 950-224, Roche) for 32 mn at 95°C for CD56 and CD133 and for 60 mn for Synaptophysine at 95°C.

Sections were incubated with:

- a mouse anti-CD56 (Novocastra, NCL-L-CD56-504) diluted at 1:100 for 32 mn. Sections were then sequentially incubated with a rabbit anti-mouse IgG2b (Abcam, ab125907) for 32 mn and then an anti-rabbit-HRP (Roche, 760-4311) for 16 mn
- a rabbit anti-CD133 (Cell signalling Technology, 86781) diluted at 1:500 for 32 minutes then an anti-rabbit-HRP (Roche, 760-4311) was applied on sections for 16 mn.
- a mouse anti-Synaptophysin (Novocastra, NCL-L-SYNAP-299) diluted at 1:200 for 32 mn. Sections were then sequentially incubated with a rabbit anti-mouse IgG1 (Abcam, ab133469) for 32 mn and then an anti-rabbit-HRP (Roche, 760-4311) for 16 mn.
- anti-Phox2B (Fine test, FNab06409) diluted at 1:200 for 20 mn on the autostainer Dako.

Staining was visualized with 3,3′-diaminobenzidine as a chromogenic substrate for 8 mn. The sections were counterstained with Gill’s Hematoxylin (Ref: 760-2021, Roche) for 8 mn and post counterstained with Bluing reagent (Ref: 760-2037, Roche) for 4 mn. The slides were washed in warm tap water with detergent and dehydrated in graded ethanol and methylcyclohexane, then coversliped in permanent mounting media Pertex (Ref: 00801-FR, HistoLab). Finally, sections were scanned with panoramic scan II (3D Histech, Budapest, Hungary) at 20X.

Five PDTs from the same culture were prepared and processed for IHC. For each PDT, we analyzed two sections along the Z-axis, resulting in 10 IHC images, numbered T1 to T10. Each pair of images (T1,T6), (T2,T7), (T3,T8), (T4,T9), and (T5,T10) comes from the same PDT.

We checked for the expression of CD133, a known stemness marker [31, 37, 39, 46] shown to suppress neuroblastoma differentiation [43] via the inhibition of RET, a tyrosine kinase receptor involved in neuroblastoma differentiation [9]. This allowed us to define two cell types: stem cells, expressing CD133, and differentiated cells expressing synaptophysin (thereafter abbreviated SYP) (see section 3.1.1).

IHCs data is available at https://osf.io/e6vug/files/osfstorage.

### 2.4 Image analysis

Images were open and analyzed using *QuPath* [2], an open source software for digital pathology image analysis.

In order to accurately identify cells in the IHC images, some parameters were modified from the default configuration when detecting positively stained cells for CD133 (respectively, SYP). We took into account the ‘background radius’, i.e. the size of the region around each cell nucleus used to estimate the local background intensity, which was set to 9 µm (resp., 6 µm).

Nuclei must have a mean intensity greater than a threshold, called intensity threshold and set at 0.05 for both markers, to be detected. Detected nuclei with an area less than a minimum, set to 2 µm for both markers, were discarded.

The mean cell count was 2,548 *±* 336 cells per section. The mean cell radius was 4.5 *±* 0.3 µm.

### 2.5 Statistical methods

To analyze the spatial distribution of stem cells in IHC images of our PDTs we focused on the following statistical indices: their percentage, Moran’s index, entropy, and centrality.

#### Percentage

It is calculated as 100 times the ratio of the number of CD133^+^ cells to the total number of cells.

#### Moran’s index

Moran’s index (thereafter named Moran’s *I*) is a statistical indicator of spatial autocorrelation, meaning it quantifies how similar or clustered values are based on their spatial locations [30]. Stem cell Moran’s *I* is computed as follows

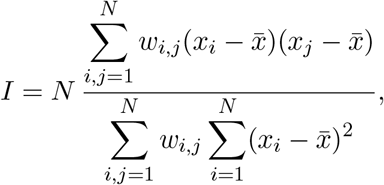

where *N* is the total number of cells, (*x*_*i*_)_*i*=1,…,*N*_ is the observation data (represented by a binary vector where *x*_*i*_ = 0 for differentiated cells and *x*_*i*_ = 1 for stem cells), 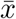 its mean, and {*w*_*i,j*_} is the neighborhood weight matrix, with *w*_*i,i*_ = 0 for all *i* and 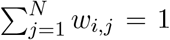. Moran’s *I* ranges between *−*1 and 1. If Moran’s *I* value is close to 0, the distribution of stem cells is random. If *I* = *−*1, the distribution is perfectly dispersed, whereas if *I* = 1 stem cells are perfectly clustered. A high Moran’s *I* indicates that stem cells tend to cluster together.

In practice, two cells are considered to be neighbors if their distance is less than or equal to 15 µm, which is in agreement with observations across all 10 IHC images.

#### Entropy

The entropy index, usually denoted by *H*, characterizes the distribution of distances between cells. The entropy index for stem cells is given by [18]

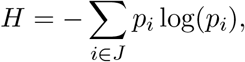

where *p*_*i*_ represents the probability that the distance between stem cells (CD133^+^) falls within bin *i* of the distance histogram, *J* being the set of all bins. This index provides complementary information about stem cell clustering.

#### Centrality

The centrality index is the ratio between the mean distance of all stem cells to the center of the IHC image of the PDT and the mean distance of all differentiated cells to the same center.

In addition, to better understand the spatial dispersion of stem cells, we computed the *intra-coefficient a*_*SS*_ introduced by Jensen and Michel [26], which quantifies the spatial dependency between cells of the same type, here the stem cell type denoted by S. More precisely,

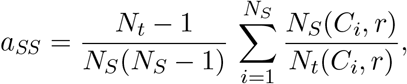

where *N*_*S*_ is the cell count of type S, *N*_*t*_ is the total number of cells, and *N*_*S*_(*C*_*i*_, *r*) (resp. *N*_*t*_(*C*_*i*_)) is the number of type S cells (resp. all types) in the neighborhood of radius *r* around stem cell *C*_*i*_ (without counting *C*_*i*_). The fraction 0*/*0 is considered equal to 1 in the right side. For all *r >* 0, the average over all cells of type *S* of this coefficient equals 1.

In [26], the analytical form of the variance of the intra-coefficient has been computed under the null hypothesis that no spatial correlation exist. Under the null hypothesis [26],

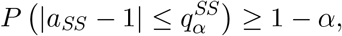

where *α* is the level of confidence and 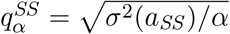.

In section 3.1.2, we consider *α* = 0.05 and compute the intra-coefficient of the IHC images along with its confidence interval 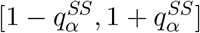, i.e. a confidence level of at least 95% under the null hypothesis. If the observed value of the intra-coefficient is greater than 1, we deduce that S-type (stem) cells tend to aggregate. Furthermore, if *a*_*SS*_ lies outside the confidence interval, i.e. 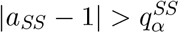, we can reject the null hypothesis.

### 2.6 Multiscale modeling framework

`Simuscale` [5] is a multiscale individual-based modeling platform for performing numerical simulations of heterogeneous populations of individual cells. It is intended for cells that experience temporal evolution (movement, differentiation, death, division) and accounts for physical and biochemical cell interactions. A `Simuscale` model is a priori described at the cellular and the population level. The cellular level describes the dynamics of single cells, and allows various formalism. The internal cell state includes, in this work, cell size and position, mRNA and protein levels for each gene (see section 2.7). At the population level, cells evolve in a bounded 3D domain, they can divide (and generate two daughter cells), differentiate, or move (see section 2.9). Spatially, cells are represented as visco-elastic spheres with a rigid core, and have two radii: the radius of the core, named ‘internal radius’, and the radius of the sphere, named ‘external radius’.

`Simuscale` implements the physical simulator that manages the simulations at the population level. Biochemical interactions occur between cells that are in contact with each other, through intercellular signals that can be known to all or to a subset of the cells only. `Simuscale` runs a simulation over a specified time interval, updating the cell population at given time steps, and it generates an output file containing the state of each cell at each time step, and the tree of cell divisions (and deaths when cells die).

### 2.7 Gene Regulatory Networks

To build the multiscale model of neuroblastoma PDTs, we first introduce a gene regulatory network (GRN), comprising the three genes CD133, SYP and Cyclin E. This GRN governs cell fate decisions through a toggle switch between CD133 and SYP, as described in section Cyclin E, a key regulator of the G1/S phase transition [25], controls the entry into proliferation. Mathematically, the GRN is written as the bursty model, a Piecewise Deterministic Markov Process (PDMP) model which couples deterministic and stochastic processes in each gene *i* ∈ *G* := {CD133, SYP, CyclinE} to produce mRNA (*M*_*i*_) and proteins (*P*_*i*_) (see [22, 23]). Furthermore, this GRN is implemented in `Simuscale`, as in [32].

More explicitly, the dynamics of mRNA and protein concentrations in a gene, (*M*_*i*_(*t*), *P*_*i*_(*t*)) for each *i* ∈ *G*, evolve deterministically with

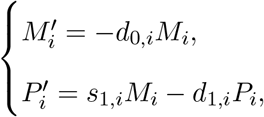

until gene *i* experiences a burst, where *s*_1,*i*_ is the protein synthesis rate of gene *i*, and *d*_0,*i*_ is the mRNA degradation rate, which is much larger than *d*_1,*i*_, the protein degradation rate. A burst causes the quantity of mRNA (*M*_*i*_) to increase by a random, discontinuous jump, sampled from an exponential distribution with a mean of 50 molecules.

We fix the degradation rate of mRNA equal to 1 h^*−*1^ for all genes, i.e. *d*_0,*i*_ = 1 h^*−*1^ for all gene *i* ∈ *G*, the protein degradation rate *d*_1,CD133_ = 0.001 h^*−*1^, *d*_1,SYP_ = *d*_1,P_ = 0.01 h^*−*1^, and *s*_1,*i*_ = 0.01 *d*_1,*i*_ h^*−*1^ for all gene *i* ∈ *G*.

Additionally, the burst rate of each gene *i* is defined as follows

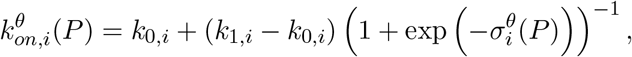

where *P* is the vector of proteins concentrations, *k*_0,*i*_ and *k*_1,*i*_ correspond to the minimum and maximum burst frequencies of gene *i*. Three scenarios are considered in this study for integrating cell-cell signaling within the burst frequency of the CD133 gene:

1. No signaling,

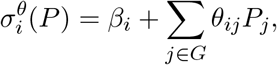

where *β*_*i*_ represents the basal activity of gene *i*, with *β*_CD133_ = *−*3, *β*_SYP_ = *β*_CyclinE_ = *−*5 (no unit). Values {*θ*_*ij*_}_*i,j*∈*G*_ denote the influence of gene *i* onto gene *j*, as encoded in the gene-gene interaction matrix between the gene protein concentrations, with their interaction strengths (no unit) specified in Table 1.
2. *S − S* signaling: Stem cell - Stem cell contact,

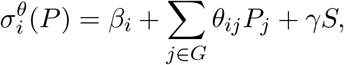

where *S* ∈ {0, 1} is the *S − S* signaling state and *γ >* 0 is the signaling strength, which will be examined in section 3.2.2.
3. signaling diffusion from stem cells,

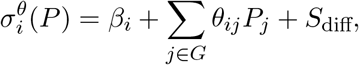

where *S*_diff_ represents the intensity of the diffusion signal perceived by a stem cell as defined in section 2.8.

### 2.8 signaling

Cell-cell contact signaling is implemented as in [32]. Stem cell signaling to other stem cells is considered in section 3.2.2, and follows this rule: two stem cells can communicate only if they are in contact (the distance of their centers is smaller than the sum of their external radii).

**Table 1:**
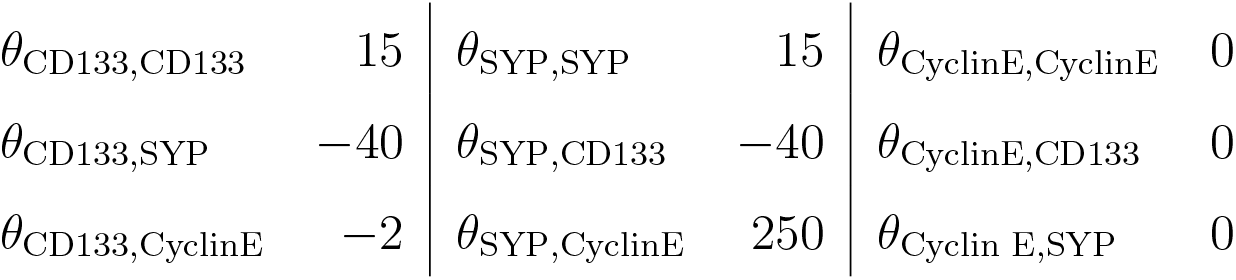
Parameter values of the gene interaction matrix.

Long-range signaling via spatially diffusing signals *S*_diff_ has recently been implemented in `Simuscale`. We assume that signals emitted by a cell (here, only a stem cell) diffuse according to a Gaussian density with variance *δ/*2, centered in a point *h* inside the cell (typically, the center of the cell). In a population of *N* cells, the total signal *S*_diff_ perceived by a cell at position *l* is the Gauss transform of *N* signal sources located at positions *h*_*j*_, with strengths *q*_*j*_,

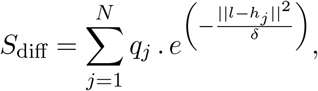

For large populations, direct computation is impractical. Instead we rely on the fast Gauss transform algorithm introduced by Greengard and Strain [20]. We fix *q*_*j*_ = 0.3 for the intensity of each stem cell signaling, and values of *δ*, which we call the *reachable area* (and is sometimes referred to as the diffusion length squared), will be studied in section 3.2.4. A movie of the diffusion signaling is available at: https://osf.io/25cy4/files/osfstorage (Diffusion_coef_2.mov).

### 2.9 Cell motion

For most scenarios considered in this work, only “mobile” movement is used for all cells [5]: in this case, cells move only because forces are applied on them, coming either from the growth of the mother cells or from daughter cells generated during cell division.

In some scenarios, “motile” cell movement (as described in [32]) is used. In this case, cells move by themselves, and we considered random cell motion within a 3D domain, where the initial velocity of each cell is set to 0.3 h^*−*1^.

When a cell encounters another cell, its velocity changes to a predefined velocity: when a stem cell encounters another stem cell, we call it a S-S velocity, when a stem cell encounters a differentiated cell (or conversely) we call it a S-D velocity, and when a differentiated cell encounters a differentiated cell we name it D-D velocity. Notably, if a stem cell simultaneously contacts multiple cell types and at least one of its neighbors is a stem cell, its velocity is set to the S-S velocity value.

### 2.10 Cell decision making

Contrary to *in vivo* tumors (see e.g. [48]), PDTs do not display a necrotic core, since gas and nutrients do flow freely through the relatively small structure. Therefore for simplification purposes, no cell death was implemented in the current version of the model.

Cell growth is a continuous linear process where cells augment their volume from *V*_birth_ = 1 (daughter cell volume at birth, radius *r* ≈ 0.62 spatial units, that is *r* ≈ 4 µm), or *V*_birth_ ≥ 1, up to *V*_division_ ≥ 2 (*r* ≈ 5 µm), where cells are ready to divide, pending upon their internal state values (see section 3.2.1). Each daughter cell inherits half the volume of the mother cell at birth, hence *V*_birth_ can be larger than 1. The conversion of a spatial domain unit in µm is based on a computed mean cell radius *r* = 0.7 corresponding to an experimentally average radius *r* = 4.5 µm.

Differentiation is also based on the cell’s internal content (see section 3.2.1). Two cell types are considered: stem cells (CD133^+^) and differentiated cells (SYP^+^). Once the cell type is acquired, some properties are directly associated to it (for example both signaling emission and perception are a stem cell characteristic) whereas other emerge from the behavior of the underlying GRN (for example, differentiated cells proliferate faster due to the strong positive influence of the SYP protein on the burst frequency of the Cyclin E gene, see section 3.2.1 for details).

### 2.11 Initial conditions

Each simulation is initialized with 15 stem cells (CD133^+^ cells) gathered side by side at the center of a 3D cube with edge length of 80 units. The spatial domain unit corresponds to 6.4 µm, so each length of the cube is 512 µm.

Molecular contents of the gene regulatory network (GRN) are initially set as follows:

- The CD133 protein value is set to twice the differentiation threshold (see section 3.2.1) value in all cells, ensuring that the first generation of daughter cells can remain stem cells. All other protein values are initialized to 0 molecule.
- mRNA values of all genes are set to 0 molecule, except for CD133, for which each cell receives an initial random value computed from an exponential burst distribution with a mean of 50 molecules.

### 2.12 Simulations

We performed 10 repetitions with the same parameter set but using different seeds, except for the diffusion study, where we performed only 5 repetitions due to computing time (calculation time exceeding 10 hours per run). All simulations were stopped when the total number of cells exceeded 50,000. This corresponds to the average cell counts from PDTs in culture when they were split, i.e. when their size approached 1 mm in diameter (see section 2.2). Interestingly, when cutting through the 3D structure, we obtained the same cell count (between 2,500 and 3,000) as observed in the IHC images (see section 2.4). All codes are available at https://github.com/ThiNhuThaoNGUYEN/SimusNeuroblas.

## 3 Results

### 3.1 Immunohistochemistry

We analyzed the 10 IHC images obtained from five PDTs from the same culture (see section 2.3). Supplementary Figure 1 shows that 100 % of the cells displayed a strong PHOX2B signal, thereby establishing their NB identity. Additionally, the amount and intensity of staining for the three antigens was fully conserved between the PDX and the PDT (and when available the original tumour), demonstrating that culture conditions did not induced any significant drift in cell identity. Finally, the expression of CD56 and synaptophysin (thereafter abbreviated SYP) was clearly less homogeneous than the expression of PHOX2B, with around 50 % of the cells stained positive. This is in line with the previous description that PDTs can recapitulate at least partly NB tissue heterogeneity.

We then focused on the spatial structure, observed in IHC images and based on the expression of CD133, defining a stem cell subpopulation, and SYP, defining a differentiated cell subpopulation.

#### 3.1.1 Spatial distribution of stem cells

Figure 1 presents the quantitative characterization of the spatial distribution of stem cells in the IHC images.

**Figure 1:**
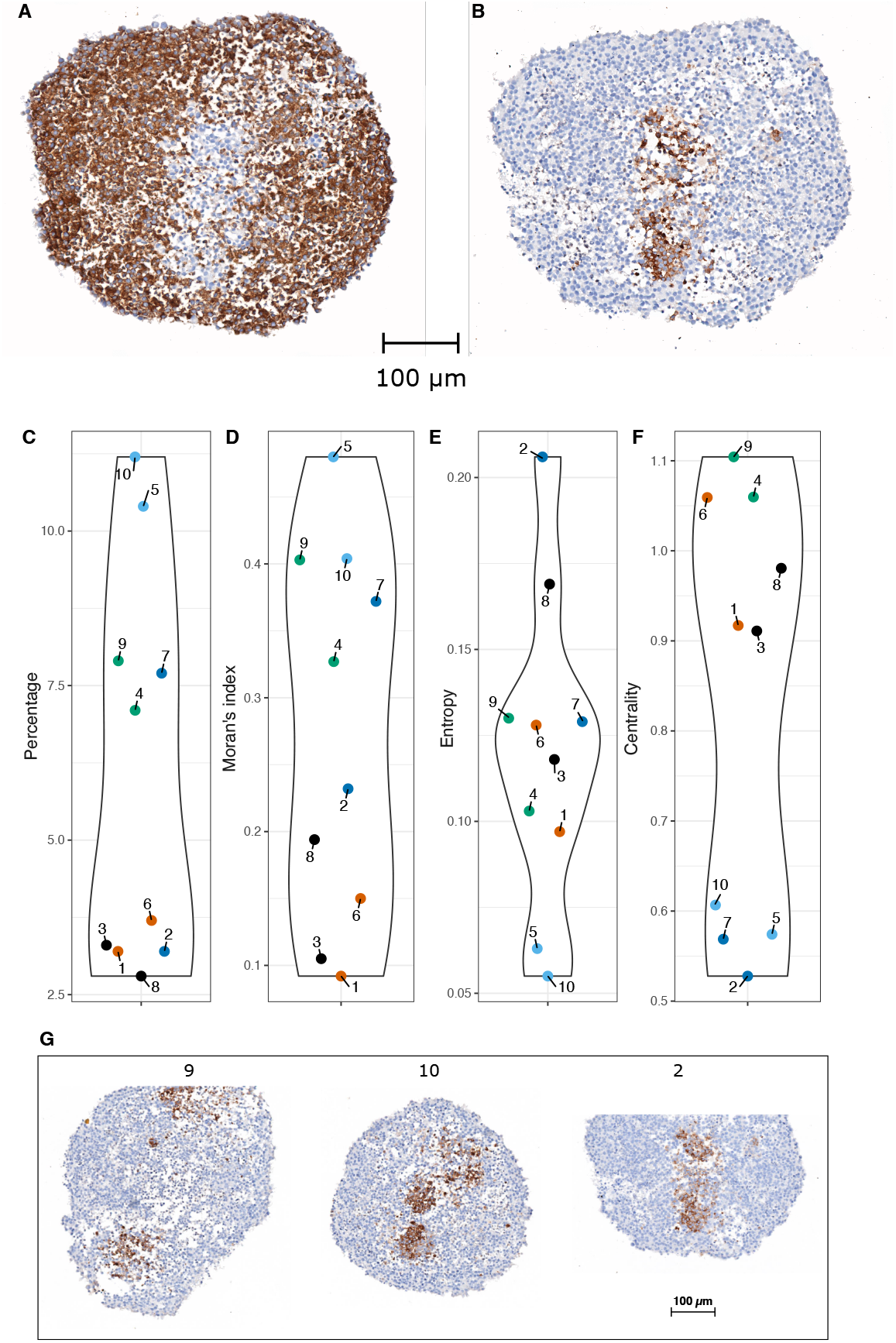
Stem cell spatial distribution. A. Percentage. B. Moran’s Index. C. Entropy. D. Centrality. Points labeled 1 to 10 correspond to the PDT images T1 to T10. The same color is used for the values computed on two sections from the same PDT. E - G. IHC images: E. CD133 labelling is shown for three PDTs. A cell appears brown if it expresses the protein; otherwise, it colors blue. F - G. Synaptophysin labelling (F) and CD133 labelling (G) are shown in two consecutive sections of the same PDT.

Figures 1A and 1B represent IHC image T7 of PDT, showing the expression of SYP and CD133, respectively, in two different sections in close proximity. It is immediately apparent that the expression of those two proteins is mutually exclusive, suggesting the existence of a toggle switch mechanism between these two genes.

We used four statistical indices (introduced in section 2.5) to quantitatively characterize the spatial distribution of CD133^+^ cells. Stem cell percentage across the 10 IHC images (Figure 1C) ranges from 2.8% to 11.2%. It is apparent that there is a relatively large variation among PDT (ranging from low values for T3 and T8 to high values for T5 and T10). Two sections arising from the same PDT display very close percentages, except for T2 and T7.

Moran’s *I* (Figure 1D) ranges from 0.09, which is close to a random distribution, to 0.48, which indicates a clear spatial clustering. Here again, some section-to-section variability from the same PDT can be observed, again more pronounced for T2 and T7.

Entropy index values (Figure 1E) range from 0.05 to 0.21. This index provides additional insights into stem cell clustering, especially in cases where Moran’s *I* values are similar, by quantifying the shape of the clustering. The more elongated the cluster is, the higher the entropy. For instance, IHC images T9 and T10 (Figure 1G) share the same Moran’s *I* value, yet their entropy values differ. This difference arises because image T9 contains two well separated stem cell clusters, whereas T10 displays more compact clusters, leading to more dispersed cell-to-cell distances.

The centrality index of stem cells (Figure 1F) shows values ranging from 0.53 to 1.1. Low centrality values indicate that stem cells are more concentrated near the center of the image, as illustrated in Figure 1E for image T2, while high centrality values highlight stem cells located away from the center of the image (see T9, on Figure 1G).

#### 3.1.2 Variance statistics for spatial dispersion of stem cells

One key question is how to ascertain the statistical significance of an empirical deviation from purely random spatial configurations. In the absence of a proper analytical expression, the significance is usually obtained by simulating many random permutation configurations by Monte Carlo simulations. When applied to Moran’s *I*, this requires a huge amount of permutations (10^7^; not shown), and therefore a very large computation time.

Jensen and Michel [26] provide an analytic formula to compute the 95% confidence interval of so-called intra-coefficient *a*_*SS*_ in the case of purely random distributions of points, and therefore alleviates the need for such permutations experiments. This allows testing *H*_0_, the hypothesis that the spatial locations of stem cells are purely random, the alternative hypothesis *H*_1_ being that cells are not randomly distributed. In this case, one has to examine whether *a*_*SS*_ stays above the reference value (i.e., 1) in which case one can conclude that cells are significantly clustered, or below the reference value in which case one can conclude that cells are significantly dispersed.

Figure 2 presents the intra-coefficient *a*_*SS*_ and its confidence interval under *H*_0_ (see section 2.5) across different values of the radius *r* for two IHC images (T1 and T4).

**Figure 2:**
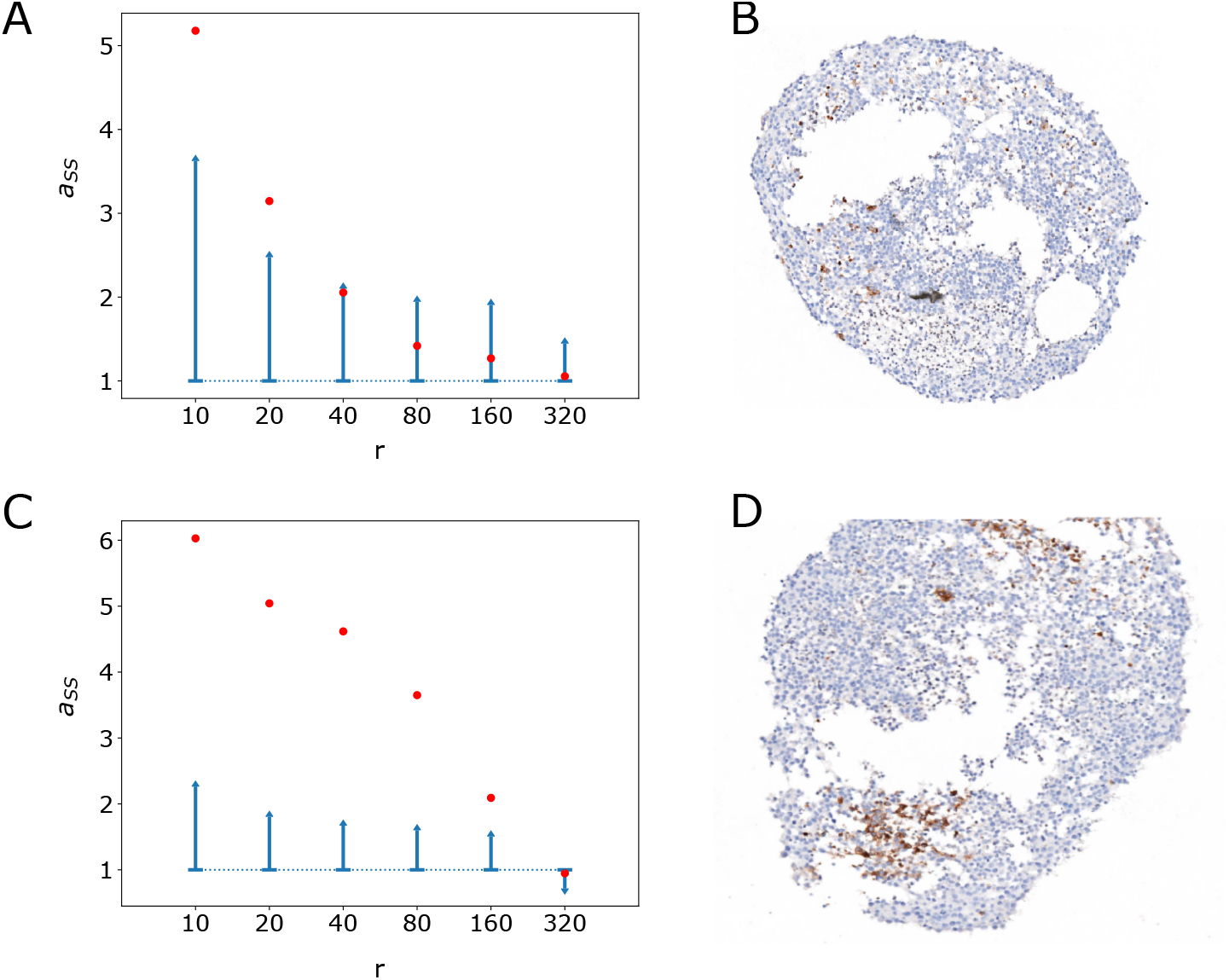
Plots of the intra-coefficient *a*_*SS*_ for stem cells with respect to the radius *r*. A and C: in red is shown the value of the intra-coefficient computed on IHC image T1 (A) and T4 (C), and in blue is shown the upper half of the 95% confidence interval, as a function of the radius (*r*, µm). The actual images of T1 and T4 are shown in B and D, respectively.

The intra-coefficient *a*_*SS*_ remained well above the confidence interval until a certain radius (40 µm for image T1 and 320 µm for image T4) is attained. This allows to reject *H*_0_ and shows that stem cells are significantly clustered up until a certain characteristic radius value. In particular, for small radius, stem cells are always clustered. For larger radius values, *a*_*SS*_ falls within the confidence interval, *H*_0_ can not be rejected, and therefore stem cells spatial locations tend to follow a random pattern.

The value of *r* at which *H*_0_ gets rejected is much higher for T4 (Figure 2C) than for T1 (Figure 2A), demonstrating that T4 displays larger clusters of stem cells than T1. It is however noteworthy that the T1 clusters that were characterized by a low Moran’s *I* nevertheless proved to be significantly different from a purely random spatial distribution. All 10 PDTs images displayed significant clustering above the expected variance under *H*_0_ (not shown, but the smallest radius value among all images was found for image T1 and equals 40 µm).

### 3.2 Computational Model

#### 3.2.1 Cell fate

We consider a dynamic model of gene activity, where molecular values are governed by a GRN that includes a toggle switch between CD133 and SYP genes (Figure 3A). These two genes allow to define two cell subpopulations: stem cells (CD133+) and differentiated cells (SYP+), as obtained experimentally (section 3.1). The GRN also incorporates the Cyclin E gene, which is used as a marker of proliferation. In this network, CD133 inhibits the burst frequency of Cyclin E, while SYP activates it, reflecting the fact that stem cells divide more slowly than differentiated cells [29]. The interaction matrix characterizing gene dynamics is provided in Table 1.

**Figure 3:**
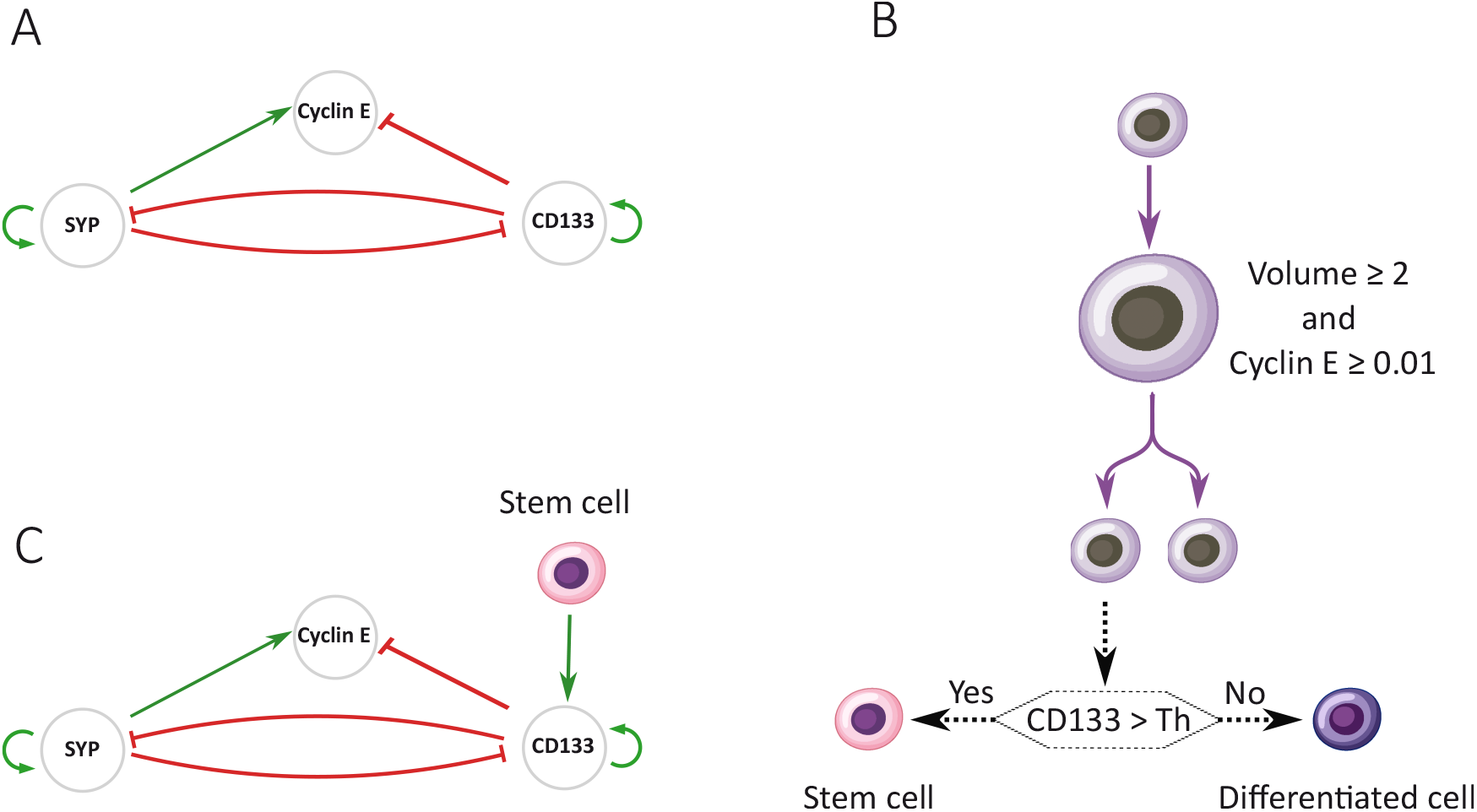
Model of cell dynamics and gene interactions. A. The first GRN implemented. It is formed of three genes, CD133, SYP, and Cyclin E, where CD133 and SYP interact through a toggle switch, and Cyclin E drives the proliferation. B. Rules for proliferation and differentiation at the cellular level. Cells grow and then divide depending on their size and the level of expression of the Cyclin E protein. The differentiation decision is taken in each daughter cells following division. The differentiation fate depends on the comparison between the CD133 protein concentration and a given threshold, Th. C. Modified GRN, where stem cells signaling influences the burst frequency of gene CD133.

We then define a model of stem and differentiated cell dynamics in which cell proliferation and differentiation dynamics are controled by the GRN (Figure 3B). Each cell has a constant volume growth rate. When its volume has at least doubled (the minimum volume is set to 1), the cell divides if the protein concentration of Cyclin E gene reaches the proliferation threshold, set to 0.01. During division, the two daughter cells inherit half of the molecular contents from their mother cell. Cell differentiation is assessed at each division: if the CD133 protein level of a daughter cell exceeds the differentiation threshold, the cell becomes or remains a stem cell; otherwise, it is deemed to be a differentiated cell.

Additionally, we propose a refined network that incorporates signaling (Figure 3C). Such signaling occurs either when a stem cell comes into contact with another stem cell, or through the diffusion of a stem cell-based signal (see section 2.8). This signaling activates CD133 by increasing its bursting frequency. As a consequence, signaling increases the likelihood of neighbouring cells to remain or become stem cells.

#### 3.2.2 Spatial structure, differentiation threshold and cell-cell contact signaling

Our objective was to generate model’s outputs ranging, for each of the four statistical indexes (see section 2.5), within the experimentally observed values.

At first, we assessed the impact of the differentiation threshold value (Figure 4A). As expected, the precentage of stem cells decreased with higher differentiation thresholds, while Moran’s *I* dropped below the lowest observed experimental values. Entropy increased with differentiation rates, reaching acceptable values when the percentage of stem cells was too small. Centrality was mostly insensitive to variations of the differentiation threshold and remained rather constant, around the value 1, therefore highlighting a lack of centrality of the stem cell population. The results showed that in the absence of any spatial rule, no noticeable spatial structure could emerge.

**Figure 4:**
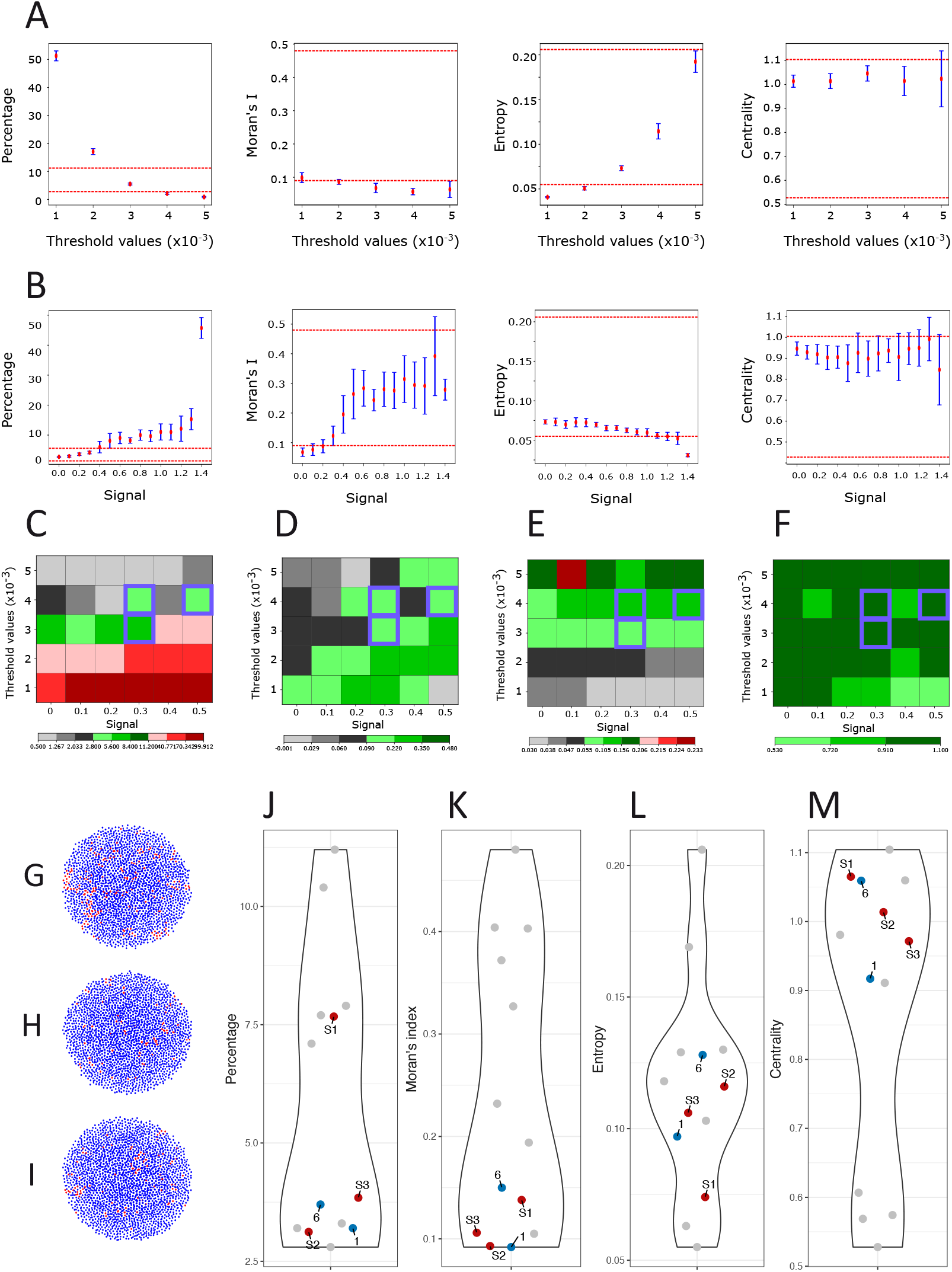
Assessing the role of differentiation threshold and cell-cell contact signaling. A. The behavior of our four indices for different differentiation threshold values. A central section was taken along the Z-axis to generate a 2D image, resembling IHC images. The mean values (red points) and standard deviations (blue lines) were then calculated. Two dashed red lines indicate the minimum and maximum values for each index based on experimental results (see Figure 1). From left to right is shown the stem cell percentage, the Moran’s *I*, the entropy index and the centrality index. B. The behavior of our four indices for different cell-cell signaling intensity, using a differentiation threshold value = 0.003. C - F. The four indices across different levels of threshold differentiation and signaling intensity (C: Percentage of stem cells; D: Moran’s *I*; E. Entropy Index; F: Centrality). Each square represents the simulated value for a specific combination of threshold differentiation (Y-axis) and signaling strength (X-axis). The color bar is from now on used to categorize the values of each index, where green indicates values within the range of minimum to maximum obtained from the experimental results (i.e. acceptable values); black indicates values below the experimental minimum; red indicates values above the experimental maximum. Each of these colors has three levels of gradation, where darker shades represent greater values. Violet squares indicate values for which all four indices are simultaneously acceptable. G - I. 2D cross section through the center of the simulated PDTs S1, S2 and S3. Stem cells are shown in red and differentiated cells in blue. J - M. Comparison of the four indices of the acceptable points S1, S2, S3 (red), and those of IHC images T1 and T6 (blue); all other images are shown in gray.

We next tested the impact of spatial signaling according to the following rule: two stem cells, when in contact, signal to each other by increasing the CD133 gene bursting rate. We set the differentiation threshold value at 0.003, a value at which the stem cell percentage, entropy, and centrality index fall within an acceptable range, even though Moran’s *I* remains below the minimum experimental value (Figure 4A). The addition of spatial signaling improved the situation, with a signaling value of 0.3 generating acceptable values for all four indices, although the Moran’s *I* value remained within the low range (Figure 4B). Increasing cell-to-cell signaling intensity increased Moran’s *I*, but pushed the percentage of stem cells index above its maximum limit.

We then assessed the joint effect of varying both the threshold differentiation and the signaling strength. For each (signal, differentiation threshold) pair, we ranked the four indices as either below, within, or above experimental range (Figure 4C-F). We obtained three pairs of parameters acceptable for all four indices: S1 = (0.3, 0.003), S2 = (0.3, 0.004), and S3 = (0.5, 0.004). The corresponding 2D sections are displayed in Figure 4G (S1), 4H (S2), and 4I (S3). The spatial distribution of stem cells is visually close to random, which is consistent with the low value of Moran’s *I* (Figure 4K). In Figures 4J - 4M, indices associated with S1, S2, and S3 are close to those of IHC images T1 and T6, which represent a case where stem cells are clustered in many small, randomly distributed groups (Figure 2B).

Altogether these results show that the addition of basic spatial rules allows at least some spatial structure to appear, although only the lower range of experimentally observed stem cell clustering can be obtained.

#### 3.2.3 Spatial structure and differential cell-cell adhesion

Cell-to-cell signaling by direct contact might be insufficient to generate the expected spatial clustering. Close examination of simulation movies indeed showed that cell growth and division break stem-to-stem contacts, which results in disrupting stem cell clusters (see an example at https://osf.io/25cy4/files/osfstorage, file Mobile.mov).

To explore how cell movement might affect clustering, we introduced an alternative movement mechanism, called “motile” in `Simuscale`. Unlike the “mobile” movement above that relies solely on passive forces from daughter cells, motile movement may allow stem cells to arrange their positions more efficiently.

We defined a default value for cell velocity of 0.3 h^*−*1^, see an example at https://osf.io/25cy4/files/osfstorage (file Motile.mov). We then made velocity contact-dependent: low velocities then mimic cell-to-cell adhesion We first assumed that both stem-to-stem (S-S) and differentiated-to-differentiated (D-D) cell adhesion would harbour the same value and that stem-to-differentiated (S-D) adhesion can somehow differ.

Figures 5A - 5D show the values of the four principal indices for different combinations of velocity parameters (S-D, S-S, D-D). A single triplet S4 = (0.256, 0, 0) h^*−*1^ was found which satisfies all four indices.

**Figure 5:**
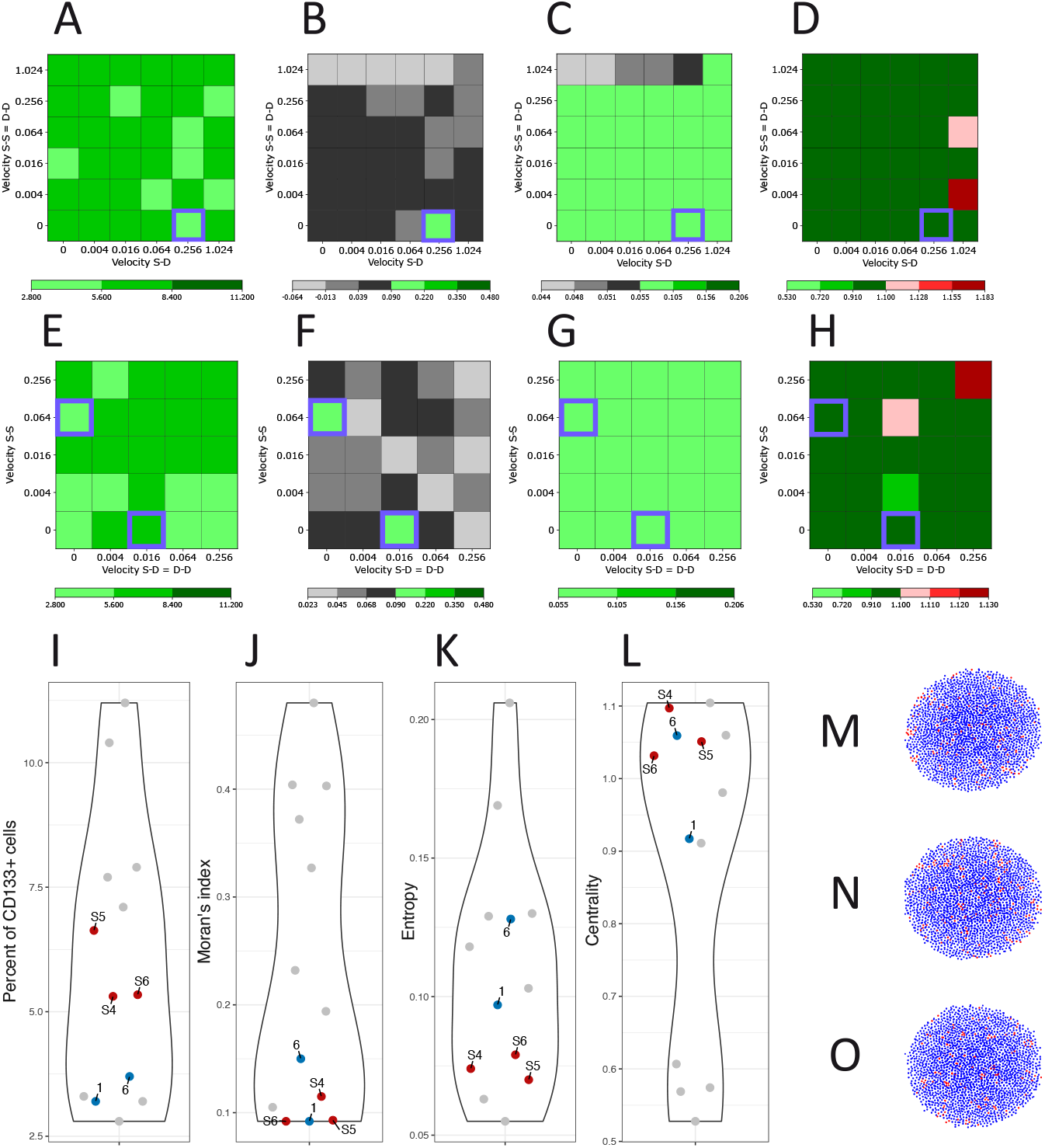
Simulation results for motile cells and differential cell-cell adhesion, while cell-cell contact signaling is reset to 0. A - D. The behavior of our four indices for different values of velocity between stem cells (S-S), equal to the velocity between differentiated cells (D-D) on the Y-axis, and the velocity between stem and differentiated cells (S-D) on the X-axis, with the acceptable points highlighted by violet squares. E - H. The four indices for different values of velocity between stem cells (S-S) on the Y-axis and between stem and differentiated cells (S-D), which is equal to the velocity between differentiated cells (D-D), on the X-axis. I - L. Comparison of the four indices of the acceptable points (red) and those of IHC images T1 and T6 (blue); all other images are shown in gray. M - O. 2D cross-section through the center of the PDT for the acceptable points.

We then examined the case where S-D adhesion between stem and differentiated cells equals the DD adhesion between differentiated cells, whereas the stem-to-stem adhesion might differ. In this case, we obtained two additional acceptable triplets of parameters, S5 = (0.016, 0, 0.016) h^*−*1^ and S6 = (0, 0.064, 0) h^*−*1^ (Figures 5E - 5H).

We also considered the case in which the adhesion between stem cells (S-S) is equal to the adhesion between stem and differentiated cells (S-D), while the adhesion between differentiated cells (D-D) may differ. However, this scenario does not provide any additional valid points (not shown).

Since Moran’s *I* of these three points was quite small, we compared all four indices associated with sections from S4, S5 and S6 with those of the IHC images. We observed that their Moran’s *I* value indeed did not differ significantly from the IHC images T1 and T6 that displayed the lower Moran’s *I* values (see Figures 5I - 5L), even though S4 to S6 displayed a higher percentage of CD133^+^ cells.

The 2D cross-sections of the PDT generated from the three configurations, S4, S5, S6, are illustrated in Figures 5M, 5N, and 5O, respectively, showing a similar spatial structure as observed in images T1 and T6 (the one with the lowest Moran’s *I*). However, we also observed that varying the cell adhesion does not significantly impact the spatial distribution of stem cells.

Altogether the addition of differential cell motility only marginally improved the ability of our modelling scheme to reproduce the experimental data.

#### 3.2.4 Role of nonlocal diffusive signaling

We finally explored a potential role for a nonlocal, spatially diffusing signal, emitted by stem cells only. The effective signaling range (from short to long range) is controled by a so-called *reachable area*, expressed as the parameter *δ*, see Section 2.8. Figure 6 depicts the simulation results when varying the reachable area.

**Figure 6:**
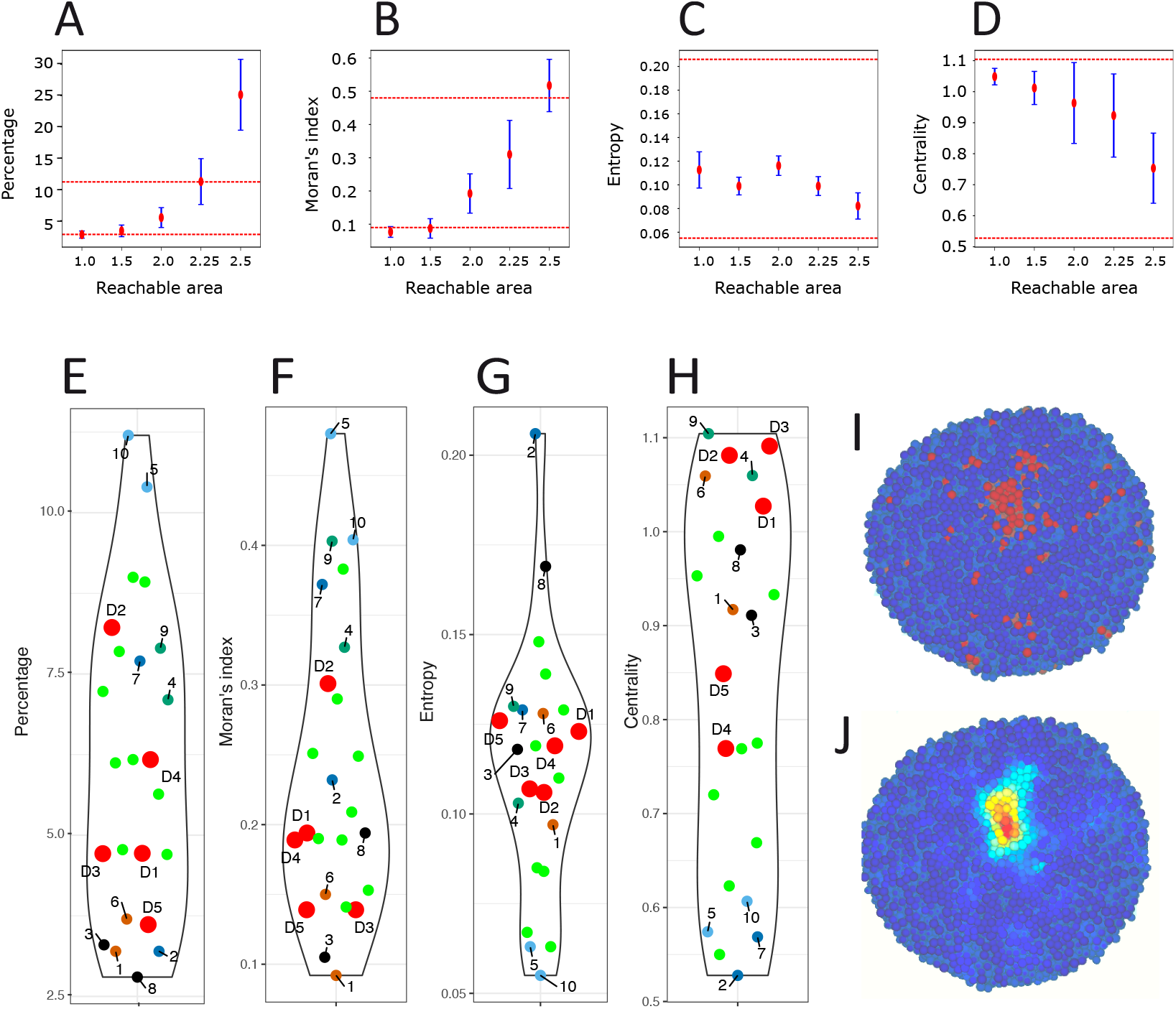
Simulation results with a spatially diffusive signal. A - D. The behavior of the four indices for different values of the reachable area *δ*. Red points represent the mean over 5 simulations, blue lines the standard deviation, and the two dashed red lines indicate the minimum and maximum experimental values for each index. E - H. Comparison of the values of the four indices between five 2D cross-sections from 5 simulations with reachable area *δ* = 2 (D1 to D5; fluorescent red), 9 successive cross-sections in one simulation (D1; fluorescent green), and the 10 IHC images from Figure 1. I. A 2D cross-section of D1 showing the spatial positioning of stem (red) and differentiated (blue) cells. J. The same 2D cross-section where the intensity of perceived cellular signaling (see section 2.8) is displayed.

As expected with a small reachable area *δ* = 1 (approximately 4.5 µm, akin to a signaling mostly felt in a one cell diameter range), the results were similar to the ones observed with direct cell-to-cell contact signaling. Also expected, when the reachable area was increased to large values (here *δ* = 2.5 being the largest tested value), the percentage of stem cells increased well beyond the maximum value observed in the experimental dataset, together with the Moran’s *I*. Indeed, the nonlocal diffusive signal maintains stem cells in their stem cell state, therefore the farther it is felt the larger the number of cells remaining stem cells.

The most interesting behavior was observed for a short-range reachable area *δ* = 2 (corresponding roughly to a uniform diffusion up to 6.4 µm), where the diffusion signaling can reach up to first-neighbor cells in any given direction. In this case, we obtained correct values for the four indices, including the highest Moran’s *I* for a correct level in the percentage of stem cells (Figure 6B).

When the simulation-based indices were plotted together with the observed experimental values (Figure 6E-H), it was clear that the simulations were able to generate quite realistic values, both in average and in their variability. Additionally, a 2D section illustrating the spatial structure of CD133^+^ cells (Figure 6I) demonstrated that the addition of short-range diffusion signal produced the most realistic stem cell spatial distributions by far.

Figure 6J shows how the signal is perceived. It demonstrates that the diffusion signal smoothened out the signaling and allowed to correct the spatial structure by reaching to stem cells that might have been separated by cellular growth.

We reasoned that at least part of the success of the short-range diffusion signal could be due to its effect on de-differentiation events. Those are events where an originally differentiated mother cell would give rise to two stem cell daughters (the level of intracellular CD133 having increased beyond the threshold during the mother lifetime). In the absence of nonlocal diffusion signaling, such de-differentiation events, though rare, occurred in 4.6 *±* 0.1% of all cell division events. When nonlocal diffusion signaling was included, de-differentiation occurred at a lower frequency, averaging 3.3 *±* 0.5% of all cell division events. This reduction could be explained by the increase in stemness associated with the wider spatial distribution of the CD133 signal, which helps stabilize the stem cell population. Thus, stem cells would be less likely to alternate between stem and differentiated phenotypes.

We finally also explored the impact of the location of the section made within the simulated PDT. For this we generated 9 successive and evenly distributed (from 10% up to 90% of the structure) cuts through the Z-plane and computed the values for the 4 indices for each section. The results (Figure 6E-H, green dots) demonstrated that there was a large variability associated to the cutting plane. For some indices, this variability is almost as large as for the 10 experimental sections.

Altogether we demonstrate that short-range spatial diffusion signaling plays a key role in successfully reconstructing the spatial structure of the PDT in this study.

## 4 Discussion

In this work, we developed a multiscale agent-based model to simulate the spatial organization of CD133^+^ cancer stem cells within neuroblastoma patient-derived tumoroids. Our results demonstrated that our *in silico* multiscale model can reproduce the main expected spatial structure both qualitatively and quantitatively, i.e. stem cells are clustered inside the PDT and surrounded by differentiated cells.

We based our study on the careful quantitative examination of immunohistochemistry (IHC) images from 5 patient-derived tumoroids grown *in vitro*. We observed the existence of two mutually exclusive gene expression patterns, with cells expressing either CD133 or SYP. This led to the definition of two phenotypes, CD133^+^ stem cell and SYP^+^ differentiated cell, and to the definition of our GRN built around a toggle switch between those two genes. The percentage of stem cells was much lower than the percentage of differentiated cells, leading to the proposed differential interaction between CD133, SYP and Cyclin E in our GRN.

A significant variability both among images originating from 2 different PDTs and between two images from the same PDT was observed. Even though all PDTs were derived from the same PDX (see section 2.1), several mechanisms have been described that can explain this variability. Although we tried hard to keep culture conditions as constant as possible, PDTs cultures proved to be quite sensitive to uncontrolled influences. For example the PDT positionning within the 96-wells plate seems to have some influence on its growth. Regarding intra-PDTs variability, the *in silico* observation of evenly spaced sections along the Z-axis clearly demonstrates that the PDT region that is retained within the cut has a strong influence on the spatial clustering indices. Ideally one would like to obtain similar results from PDTs grown *in vitro*, i.e. estimate the value of our four indices on many consecutive cuts. Preliminary results (not shown) suggest that this might be feasible, but with much less control on the positioning of the cut as compared to *in silico* experiments. Furthermore, technical artefacts during the cut could also limit the extent of section-to-section reproducibility. An elegant alternative would consist in using Light sheet fluorescence microscopy which allows to quickly observe and quantitatively evaluate biological processes and structures in three dimensions with minimal light-induced damage and fading [40]. This should results in 3D images on which we could compute spatial statistics directly comparable to our 3D *in silico* simulations.

In the present work we did not explore the role of initial seeding conditions. We seeded only a fixed number of stem cells, while in culture the initial cell composition can fluctuate. It could therefore be interesting to explore more diverse initial conditions, by varying both the number and the type of seeded cells. This would allow us to assess how the initial cell composition influences the resulting spatial organization and dynamics of the PDT. The results could then be compared to PDTs originating from FACS-sorted stem or differentiated cells or a mix of both.

One remaining question concerns the molecular nature of the diffusive signal. A quick survey of the litterature regarding autocrine signaling in cancer stem cells reveals multiple possible candidates, although none were yet identified in neuroblastomas: for example, an autocrine loop using the vascular endothelial growth factor-C (VEGF-C) was described for various cancers [47] or an autocrine loop using the Hepatocyte growth factor (HGF)/c-Met pathway was described for Renal cell carcinoma cancer stem cells [38]. Furthermore, TGF-*β* has been demonstrated to promote stem cell properties through well documented autocrine loops [11], as well as inducing and maintaining epithelial-mesenchymal transition in transformed human mammary epithelial cells by an autocrine signaling loop, in conjonction with the Wnt signaling pathway [10]. In this last case, the authors have shown that the inhibition of these pathways is sufficient to prevent the acquisition of stem properties by cells.

We could test various ligand-receptor coexpression in our cancer stem cells (CSC) using single cell transcriptomics data that are currently being generated on our PDTs, in order to isolate putative autocrine signaling mechanisms. This could lead to propose a therapeutic strategy in line with [46] that proposed direct targeting of CSCs, using various drugs like AZD9150, a STAT3-targetting agent, or epigenetic drugs such as EZH2 or SETD8 inhibitors to push CSCs out of their stem-like, drug-resistant state, or retinoic acid to force CSC differentiation.

The present framework could be extended to model *bona fide* neuroblastoma tumors. A possible source for experimental evidence could come from recently published spatial transcriptomics studies [35]. One of the main difficulty that can be envisionned is to correctly define the complexity level at which the model should be defined. An inspiration could come from the work of Borau et al. [8] who presented the first multiscale orchestrated framework that integrates molecular, cellular, and tissue-level processes involved in neuroblastoma, using patient-specific data. Major differences with this work would consist in introducing a specific CSC compartment and in using an explicit dynamical GRN to produce the cell’s molecular content. We are currently working on the inference of such a more realistic GRN using the CARDAMOM inference tool [45]. Such an approach could then lead to generalize the framework to solid tumors when spatial transcriptomics data are available.

## Acknowledgments

This work was supported by the PEPR Santé Numérique under Project AI4scMED MultiScale AI for Single Cell-Based Precision Medicine (ANR-22-PESN-0002) and by the Ligue Contre le Cancer (comités du Rhône et de l’Ardèche)

We would like to acknowledge the invaluable support of Julie Valantin, Nicolas Gadot and Elodie Voilin from the Research Pathology Platform East for their help with IHC; the Service de Biopathologie du Centre Leon Berard for the realization of Phox2B IHC; and Isabelle Goddard from the PPAC: Plateforme du Petit Animal du CRCL for her help with mice experiments.

We thank Pablo Jensen for insightful discussions regarding the computation of the *intra-coefficient a*_*SS*_.

We also thank the PSMN and the CC-IN2P3 for their generous access to computational resources.

## Author Contributions

Conceptualization, T.N.T.N, O.G and F.C; Data Generation, C.K., E.V., L.B., O.G. and S.G.-G Funding acquisition, O.G and F.C; Investigation, T.N.T.N, O.G and F.C; Methodology, T.N.T.N, O.G and F.C; Project administration, O.G and F.C; Software, T.N.T.N, S.B; Supervision, O.G and F.C; Visualization, T.N.T.N, O.G and F.C; Writing – original draft, T.N.T.N, O.G, and F.C; Writing – review & editing, All.

## Supplementary Figures

**Supplementary Figure 1:**
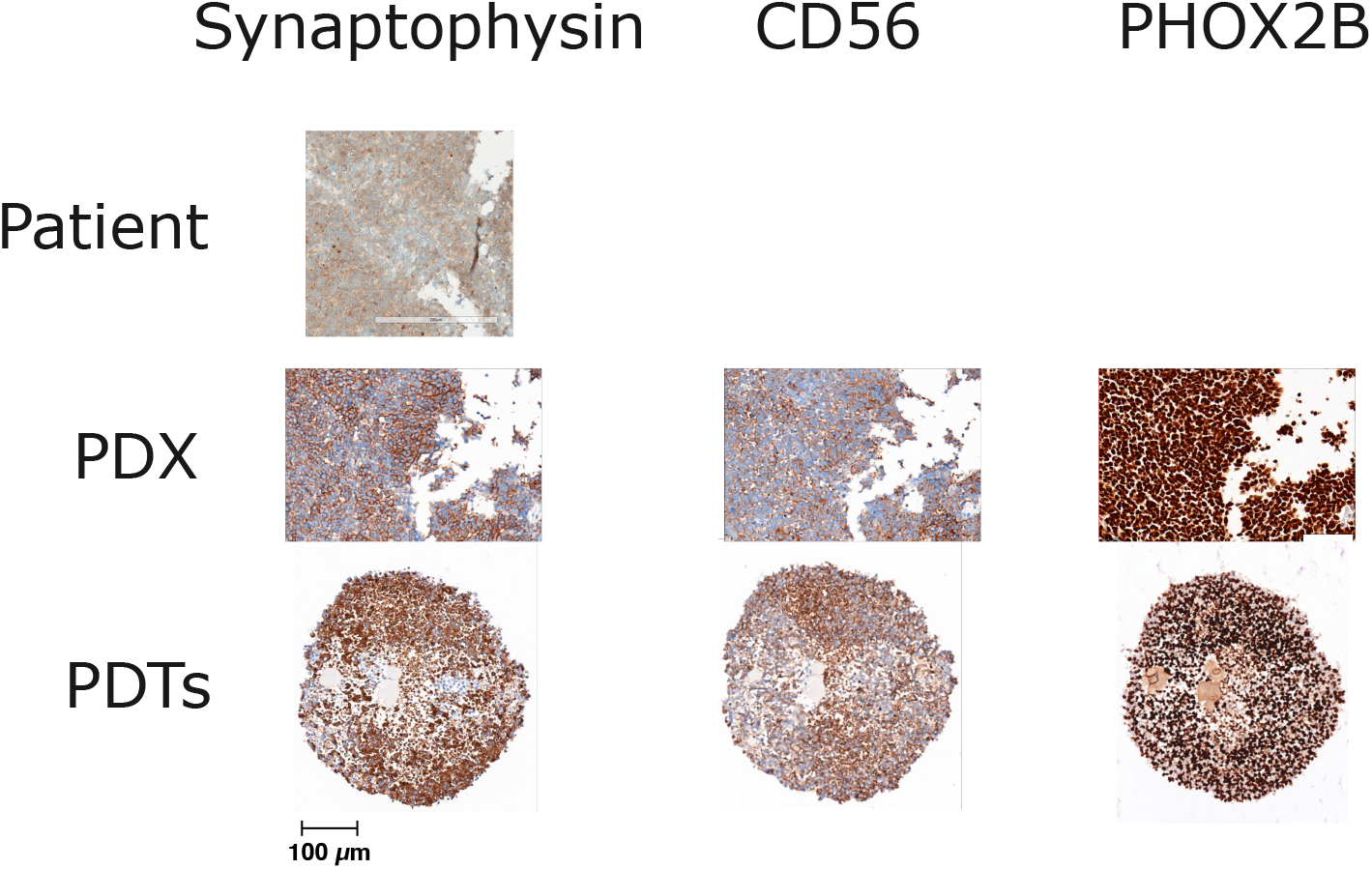
IHC staining using clinical markers routinely used for NB diagnosis. Shown is the original tumor (image courtesy of the St Jude Hospital, first line), our own derived PDX (second line), and the corresponding derived PDT (third line) at passage 10. Positive cells are stained in brown.

## References

[1] Almstedt, E., Elgendy, R., Hekmati, N., Rosén, E., Wärn, C., Olsen, T. K., Dyberg, C., Doroszko, M., Larsson, I., Sundström, A., Arsenian Henriksson, M., Påhlman, S., Bexell, D., Vanlandewijck, M., Kogner, P., Jörnsten, R., Krona, C., and Nelander, S. (2020). Integrative discovery of treatments for high-risk neuroblastoma. Nature Communications, 11(1):71.

[2] Bankhead, P., Loughrey, M. B., Fernandez, J. A., Dombrowski, Y., McArt, D. G., Dunne, P. D., McQuaid, S., Gray, R. T., Murray, L. J., Coleman, H. G., James, J. A., Salto-Tellez, M., and Hamilton, P. W. (2017). Qupath: Open source software for digital pathology image analysis. Sci Rep, 7(1):16878.

[3] Barberis, L. (2021). Radial percolation reveals that cancer stem cells are trapped in the core of colonies. Papers in Physics, 13:130002.

[4] Bate-Eya, L. T., Ebus, M. E., Koster, J., den Hartog, I. J., Zwijnenburg, D. A., Schild, L., van der Ploeg, I., Dolman, M. E., Caron, H. N., Versteeg, R., and Molenaar, J. J. (2014). Newly-derived neuroblastoma cell lines propagated in serum-free media recapitulate the genotype and phenotype of primary neuroblastoma tumours. Eur J Cancer, 50(3):628–37.

[5] Bernard, S., Crauste, F., Gandrillon, O., Knibbe, C., and Parsons, D. (2024). Simuscale: A modular framework for multiscale single-cell modelling. Technical Report RT-0520, Inria Lyon.

[6] Biava, P. M., Basevi, M., Biggiero, L., Borgonovo, A., Borgonovo, E., and Burigana, F. (2011). Cancer cell reprogramming: stem cell differentiation stage factors and an agent based model to optimize cancer treatment. Curr Pharm Biotechnol, 12(2):231–42.

[7] Bohrer, C. H. and Larson, D. R. (2021). The stochastic genome and its role in gene expression. Cold Spring Harb Perspect Biol.

[8] Borau, C., Wertheim, K. Y., Hervas-Raluy, S., Sainz-DeMena, D., Walker, D., Chisholm, R., Richmond, P., Varella, V., Viceconti, M., Montero, A., Gregori-Puigjane, E., Mestres, J., Kasztelnik, M., and Garcia-Aznar, J. M. (2023). A multiscale orchestrated computational framework to reveal emergent phenomena in neuroblastoma. Comput Methods Programs Biomed, 241:107742.

[9] Cerchia, L., D’Alessio, A., Amabile, G., Duconge, F., Pestourie, C., Tavitian, B., Libri, D., and de Franciscis, V. (2006). An autocrine loop involving ret and glial cell-derived neurotrophic factor mediates retinoic acid-induced neuroblastoma cell differentiation. Mol Cancer Res, 4(7):481–8.

[10] Dalla Pozza, E., Forciniti, S., Palmieri, M., and Dando, I. (2018). Secreted molecules inducing epithelial-to-mesenchymal transition in cancer development. Seminars in Cell & Developmental Biology, 78:62–72.

[11] Derynck, R., Turley, S. J., and Akhurst, R. J. (2021). Tgfβ biology in cancer progression and immunotherapy. Nat Rev Clin Oncol, 18(1):9–34.

[12] Enderling, H. (2015). Cancer stem cells: small subpopulation or evolving fraction? Integrative Biology, 7(1):14–23.

[13] Enderling, H., Anderson, A. R., Chaplain, M. A., Beheshti, A., Hlatky, L., and Hahnfeldt, P. (2009a). Paradoxical dependencies of tumor dormancy and progression on basic cell kinetics. Cancer Res, 69(22):8814–21.

[14] Enderling, H., Hlatky, L., and Hahnfeldt, P. (2009b). Migration rules: tumours are conglomerates of self-metastases. Br J Cancer, 100(12):1917–25.

[15] Enderling, H., Hlatky, L., and Hahnfeldt, P. (2010). The promoting role of a tumour-secreted chemorepellent in self-metastatic tumour progression. Mathematical medicine and biology : a journal of the IMA, 29:21–9.

[16] Fotinós, J., Marks, M. P., Barberis, L., and Vellón, L. (2024). Assessing the distribution of cancer stem cells in tumorspheres. Scientific Reports, 14(1):11013.

[17] Fusco, P., Parisatto, B., Rampazzo, E., Persano, L., Frasson, C., Di Meglio, A., Leslz, A., Santoro, L., Cafferata, B., Zin, A., Cimetta, E., Basso, G., Esposito, M. R., and Tonini, G. P. (2019). Patient-derived organoids (pdos) as a novel in vitro model for neuroblastoma tumours. BMC Cancer, 19(1):970.

[18] Gandrillon, O., Gaillard, M., Espinasse, T., Garnier, N. B., Dussiau, C., Kosmider, O., and Sujobert, P. (2021). Entropy as a measure of variability and stemness in single-cell transcriptomics. Current Opinion in Systems Biology, 27:100348.

[19] Gonzalez, A. L., Luciana, L., Le Neve, C., Valantin, J., Francols, L., Gadot, N., Vanbelle, C., Davignon, L., and Broutier, L. (2021). Staining and high-resolution imaging of three-dimensional organoid and spheroid models. Journal of visualized experiments : JoVE, 169:e62280.

[20] Greengard, L. and Strain, J. (1991). The fast gauss transform. SIAM Journal on Scientific and Statistical Computing, 12(1):79–94.

[21] Hagolani, P. F., Zimm, R., Marin-Riera, M., and Salazar-Ciudad, I. (2019). Cell signaling stabilizes morphogenesis against noise. Development, 146(20).

[22] Herbach, U. (2023). Harissa: Stochastic simulation and inference of gene regulatory networks based on transcriptional bursting. In Pang, J. and Niehren, J., editors, Computational Methods in Systems Biology, pages 97–105. Springer Nature Switzerland.

[23] Herbach, U., Bonnaffoux, A., Espinasse, T., and Gandrillon, O. (2017). Inferring gene regulatory networks from single-cell data: a mechanistic approach. BMC Systems Biology, 11:105.

[24] Hillen, T., Enderling, H., and Hahnfeldt, P. (2013). The tumor growth paradox and immune system-mediated selection for cancer stem cells. Bulletin of Mathematical Biology, 75(1):161– 184.

[25] Hwang, H. C. and Clurman, B. E. (2005). Cyclin e in normal and neoplastic cell cycles. Oncogene, 24(17):2776–86.

[26] Jensen, P. and Michel, J. (2011). Measuring spatial dispersion: exact results on the variance of random spatial distributions. The Annals of Regional Science, 47(1):81–110.

[27] Luksch, R., Castellani, M. R., Collini, P., De Bernardi, B., Conte, M., Gambini, C., Gandola, L., Garaventa, A., Biasoni, D., Podda, M., Sementa, A. R., Gatta, G., and Tonini, G. P. (2016). Neuroblastoma (peripheral neuroblastic tumours). Crit Rev Oncol Hematol, 107:163–181.

[28] M. Kholosy, W., Derieppe, M., van den Ham, F., Ober, K., Su, Y., Custers, L., Schild, L., M. J. van Zogchel, L., M. Wellens, L., R. Ariese, H., Szanto, C. L., Wienke, J., Dierselhuis, M. P., van Vuurden, D., Dolman, E. M., and Molenaar, J. J. (2021). Neuroblastoma and dipg organoid coculture system for personalized assessment of novel anticancer immunotherapies. Journal of Personalized Medicine, 11(9).

[29] Moore, N. and Lyle, S. (2011). Quiescent, slow-cycling stem cell populations in cancer: A review of the evidence and discussion of significance. Journal of Oncology, 2011:396076.

[30] Moran, P. A. P. (1950). Notes on continuous stochastic phenomena. Biometrika, 37(1-2):17–23.

[31] Moreno-Londono, A. P. and Robles-Flores, M. (2024). Functional roles of cd133: More than stemness associated factor regulated by the microenvironment. Stem Cell Rev Rep, 20(1):25– 51.

[32] Nguyen, T. N. T., Martin, M., Arpin, C., Bernard, S., Gandrillon, O., and Crauste, F. (2024). In silico modelling of CD8 T cell immune response links genetic regulation to population dynamics. ImmunoInformatics, 15.

[33] Noble, R., Tasaki, K., Noble, P. J., and Noble, D. (2019). Biological relativity requires circular causality but not symmetry of causation: So, where, what and when are the boundaries? Frontiers in Physiology, 10:827.

[34] Padovan-Merhar, O. M., Raman, P., Ostrovnaya, I., Kalletla, K., Rubnitz, K. R., Sanford, E. M., Ali, S. M., Miller, V. A., Mossé, Y. P., Granger, M. P., Weiss, B., Maris, J. M., and Modak, S. (2016). Enrichment of targetable mutations in the relapsed neuroblastoma genome. PLOS Genetics, 12:1–13.

[35] Patel, A. G., Ashenberg, O., Collins, N. B., Segerstolpe, A., Jiang, S., Slyper, M., Huang, X., Caraccio, C., Jin, H., Sheppard, H., Xu, K., Chang, T.-C., Orr, B. A., Shirinifard, A., Chapple, R. H., Shen, A., Clay, M. R., Tatevossian, R. G., Reilly, C., Patel, J., Lupo, M., Cline, C., Dionne, D., Porter, C. B. M., Waldman, J., Bai, Y., Zhu, B., Barrera, I., Murray, E., Vigneau, S., Napolitano, S., Wakiro, I., Wu, J., Grimaldi, G., Dellostritto, L., Helvie, K., Rotem, A., Lako, A., Cullen, N., Pfaff, K. L., Karlström, A., Jane-Valbuena, J., Todres, E., Thorner, A., Geeleher, P., Rodig, S. J., Zhou, X., Stewart, E., Johnson, B. E., Wu, G., Chen, F., Yu, J., Goltsev, Y., Nolan, G. P., Rozenblatt-Rosen, O., Regev, A., and Dyer, M. A. (2024). A spatial cell atlas of neuroblastoma reveals developmental, epigenetic and spatial axis of tumor heterogeneity. bioRxiv, page 2024.01.07.574538.

[36] Reid, M. A., Dai, Z., and Locasale, J. W. (2017). The impact of cellular metabolism on chromatin dynamics and epigenetics. Nat Cell Biol, 19(11):1298–1306.

[37] Ross, R. A., Walton, J. D., Han, D., Guo, H. F., and Cheung, N. K. (2015). A distinct gene expression signature characterizes human neuroblastoma cancer stem cells. Stem Cell Res, 15(2):419–26.

[38] Silva Paiva, R., Gomes, I., Casimiro, S., Fernandes, I., and Costa, L. (2020). c-met expression in renal cell carcinoma with bone metastases. J Bone Oncol, 25:100315.

[39] Singh, S. K., Clarke, I. D., Hide, T., and Dirks, P. B. (2004). Cancer stem cells in nervous system tumors. Oncogene, 23(43):7267–73.

[40] Stelzer, E. H. K., Strobl, F., Chang, B.-J., Preusser, F., Preibisch, S., McDole, K., and Fiolka, R. (2021). Light sheet fluorescence microscopy. Nature Reviews Methods Primers, 1(1):73.

[41] Stewart, E., Federico, S. M., Chen, X., Shelat, A. A., Bradley, C., Gordon, B., Karlstrom, A., Twarog, N. R., Clay, M. R., Bahrami, A., Freeman, B. B., r., Xu, B., Zhou, X., Wu, J., Honnell, V., Ocarz, M., Blankenship, K., Dapper, J., Mardis, E. R., Wilson, R. K., Downing, J., Zhang, J., Easton, J., Pappo, A., and Dyer, M. A. (2017). Orthotopic patient-derived xenografts of paediatric solid tumours. Nature, 549(7670):96–100.

[42] Stiehl, T., Baran, N., Ho, A. D., and Marciniak-Czochra, A. (2015). Cell division patterns in acute myeloid leukemia stem-like cells determine clinical course: a model to predict patient survival. Cancer Res, 75(6):940–9.

[43] Takenobu, H., Shimozato, O., Nakamura, T., Ochiai, H., Yamaguchi, Y., Ohira, M., Nakagawara, A., and Kamijo, T. (2011). Cd133 suppresses neuroblastoma cell differentiation via signal pathway modification. Oncogene, 30(1):97–105.

[44] Vegliante, R., Pastushenko, I., and Blanpain, C. (2022). Deciphering functional tumor states at single-cell resolution. The EMBO Journal, 41(2):e109221.

[45] Ventre, E., Herbach, U., Espinasse, T., Benoit, G., and Gandrillon, O. (2023). One model fits all: combining inference and simulation of gene regulatory networks. PLoS Comput Biol, 19(3):e1010962.

[46] Veschi, V., Verona, F., and Thiele, C. J. (2019). Cancer stem cells and neuroblastoma: Characteristics and therapeutic targeting options. Front Endocrinol (Lausanne), 10:782.

[47] Wang, C. A. and Tsai, S. J. (2015). The non-canonical role of vascular endothelial growth factor-c axis in cancer progression. Exp Biol Med (Maywood), 240(6):718–24.

[48] Wang, Z., Butner, J. D., Kerketta, R., Cristini, V., and Deisboeck, T. S. (2015). Simulating cancer growth with multiscale agent-based modeling. Seminars in Cancer Biology, 30:70–78.

